# Evidence of complete myofibril remodeling after severe damage in adult *Drosophila*

**DOI:** 10.64898/2026.09.10.750754

**Authors:** Tiara Mulder, Kate MacKee, Santiago Castillo-Ramírez, Francesca Di Cara, Nicanor González-Morales

## Abstract

Muscle function depends on the ability of myofibrils to withstand and repair mechanical damage, yet how adult muscles remodel damaged myofibrils remains poorly understood. Here, we establish a *Drosophila* model that allows the induction and longitudinal visualization of extensive myofibril damage and recovery in intact adult femur muscles. Sustained muscle depolarization caused extensive disruption of myofibrillar organization, with severe damage characterized by near-complete loss of Z-disc structures. Remarkably, myofibril architecture and muscle function were largely restored within days, revealing a substantial capacity for myofibril reconstruction in adult femur muscles. We identified two distinct states of myofibril damage, mild and severe, with mild damage appearing before severe damage during aging, suggesting a progressive process of myofibril deterioration and repair failure. We further show that filamins mechanosignaling is required for efficient myofibril remodeling. Following damage, wild-type filamin redistributes from the Z-disc and accumulates outside the myofibrils, whereas constitutively open filamin remains Z-disc-associated and constitutively closed filamin redistributes but results in increased damage and impaired recovery. These findings suggest that effective repair requires dynamic transitions between filamin conformational states and that filamin redistribution is an active component of the damage response rather than simply a consequence of muscle injury. During aging, filamin progressively redistributes from the Z-disc and muscle damage accumulates, with severe damage increasing after the appearance of mild damage. Together, our findings reveal a previously unappreciated capacity of adult muscle to reassemble damaged myofibrils and identify filamin mechanosignaling as a key component of this repair process, providing a framework for understanding how defective mechanosensing may contribute to age-related muscle decline and muscle disease.

## Introduction

Muscles are multinucleated cells; their cytoskeleton is composed of long cables or myofibrils made up of serially repeated sarcomeres, which in turn are composed of antiparallel thin and thick filaments. Thin filaments anchor to a region of the sarcomere called the Z-disc, while the thick filaments are aligned at the center of a sarcomere in a region called the M-line (Schöck & González-Morales, 2022).

Muscles frequently sustain mechanical damage which are efficiently repaired (Højfeldt et al., 2025). Severe muscle injuries that result in myofiber necrosis are repaired through regenerative myogenesis involving activation, proliferation, and differentiation of muscle satellite cells (Højfeldt et al., 2025). In contrast, localized myofibrillar damage caused by strong muscle contractions, can be repaired within the existing myofiber through a cell-autonomous mechanism that does not require satellite-cells (Roman et al., 2021). In vertebrates, directly after myofibril damage, the proteins filamin-C and Hsp27 accumulate at the damage sites (Roman et al., 2021).

In insects, evidence of muscle repair at the adult stage is scarce. A limited number of sarcomere proteins undergo turnover in adult myofibrils, but they are required to prevent early degeneration of muscles (Perkins & Tanentzapf, 2014). In adult *Drosophila*, the indirect flight muscles (IFM) are resistant to puncture mechanical damage (Chaturvedi et al., 2017), these ability comes from the action of a subset of hemocyte cells positive for Zfh1 and Pxn that remodel the extracellular matrix and contribute to structural stabilization of injured muscle (Ammar et al., 2026; Boukhatmi & Bray, 2018). The Colorado potato beetle (*L.decemlineata*) degrades muscle mitochondria through a parkin dependant pathway during diapause and makes them again after diapause (Lebenzon et al., 2022), suggesting that adult stage insects can dynamically remodel muscles subcellular structures.

Additional evidence of an adult muscle repair system in insects comes from functional studies of filamin, which is continuously expressed in adult muscles and is required for myofibril organization (González-Morales et al., 2017; Spletter et al., 2018). Filamin is localized at the Z-disc, where it connects adjacent sarcomeres (González-Morales & Schöck, 2020; Szikora et al., 2019). Filamin senses mechanical forces through its mechanosensory region (MSR), which adopts an open conformation under tension, exposing protein-binding sites that are masked in the closed state (Lad et al., 2007, 2008; Pentikäinen & Ylänne, 2009; Pudas et al., 2005; Rognoni et al., 2012). Biochemical and computational screens suggest many proteins bind specifically to the open MSR (Johnson & González-Morales, 2025; Korkiamäki et al., 2026). Mechanically damaged MSR is detected by the conserved kinase NUAK and degraded (Brooks et al., 2020).

Without a functional MSR, myofibrils break but only upon contractile forces, suggesting that like in vertebrates filamin is essential for muscle maintenance (Fisher et al., 2024; Leber et al., 2016; Mulder et al., 2025).

Here we test the presence of myofibril remodelling after damage using the *Drosophila* femur muscles in combination with optogenetic and thermogenetic depolarization. The femur is primarily composed of the tibia levator and depressor muscles, which originate from a subpopulation of Twist-expressing cells in the leg imaginal disc (Guillermin et al., 2026; Soler et al., 2004). Unlike other insect muscles, femur muscles are organized around internal tendons and are innervated by motor neurons that regulate their activation and force output (Baek & Mann, 2009; Brierley et al., 2012; Lesser et al., 2024). The femur muscles are tubular and synchronous, with branched, cross-striated myofibrils and mitochondria form a grid-like pattern embedded within leg muscle myofibrils (Ajayi et al., 2022; Avellaneda et al., 2021). Despite these differences, both muscle types share core sarcomere proteins, including Zasp52, Zasp66, α-actinin, actin, myosin heavy chain, and obscurin (González-Morales et al., 2019; Katzemich et al., 2012; Sarov et al., 2016).

## Results

### Depolarization-induced muscle damage leads to complete myofibril loss

We focused on the femur levator muscles because they are highly active during locomotion, express high levels of filamin (Fig. 1A and B), and can be imaged through the transparent cuticle without dissection. To induce controlled muscle damage, we expressed the red-shifted channelrhodopsin CsChrimson in muscles using Mef2-Gal4. Red-light illumination induced rapid paralysis that was immediately reversible after cessation of stimulation (Fig. 1C and Supplementary Fig. 1). A 2-hrs exposure was sufficient to induce severe muscle dysfunction without lethality and was therefore used for subsequent experiments (Supplementary Fig. 2A-C). Following 2 hrs of red-light stimulation, Mef2>CsChrimson flies exhibited a severe reduction in locomotor performance that progressively recovered over seven days, returning to pre-stimulation levels (Fig. 1D). To determine whether this response was specific to optogenetic stimulation, we repeated the experiment using the temperature-sensitive cation channel TrpA1 (Hamada et al., 2008). Muscle activation at 37°C produced a similar paralysis and progressive recovery of climbing ability (Fig. 1C and E). Muscle birefringence was lost after depolarization but recovered within 7 days, confirming that sustained muscle depolarization induces severe but reversible damage at the cellular level. (Fig. 1F).

**Figure 1.**
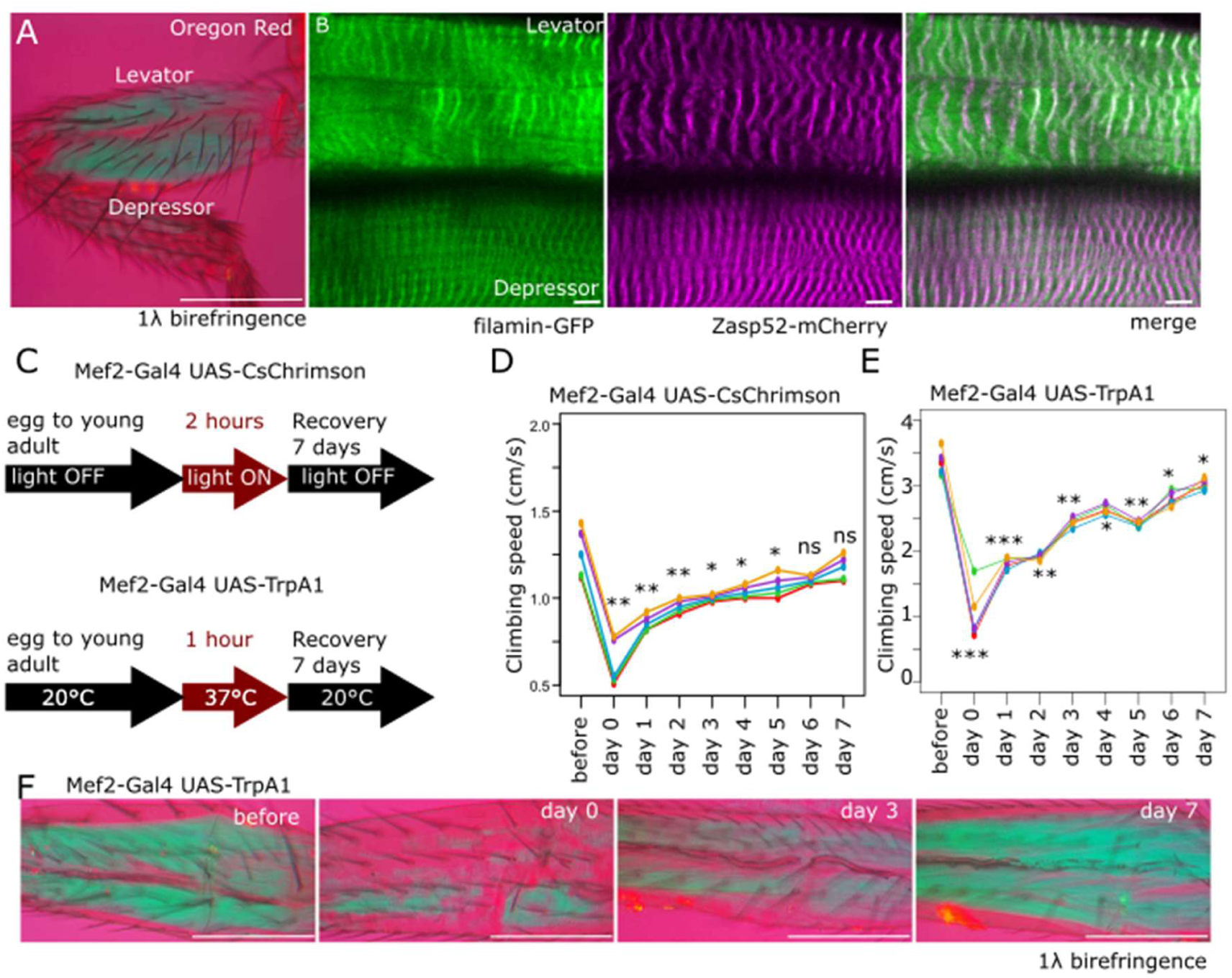
Drosophila femur muscles and depolarization-induced muscle damage. A) Polarization microscopy image of a Drosophila leg acquired using a 25× objective. Well-oriented muscle fibers exhibit bright blue birefringence against the magenta background produced by a first-order red retardation plate. B) Representative confocal image of femur muscle expressing filamin-WT-GFP and Zasp52-mCherry. Filamin is diffusely distributed at the Z-disc, whereas Zasp52 is predominantly restricted to the Z-disc. C) Schematic representation of the light-and temperature-induced muscle depolarization protocols. D) Climbing speed of Mef2-Gal4 > UAS-CsChrimson-Venus flies before and after depolarization-induced damage and during the subsequent progressive recovery period. E) Climbing speed of Mef2-Gal4 > UAS-TrpA1 flies before and after depolarization-induced damage, showing a similar progressive recovery. F) Polarization microscopy images of femur muscles during damage and recovery. Scale bars: A, 5 µm; B, 70 µm.

### Myofibrils fully reassemble after a few days of myofibril disintegration

To determine the structural basis of this functional impairment, we visualized femur muscles expressing Zasp52-mCherry and classified damaged regions as mild, in which Z-disc organization was disrupted but still detectable, or severe, in which Z-disc organization was completely lost. In Mef2>CsChrimson muscles, severe damage occupied approximately 60% of the muscle area immediately after stimulation but was absent after three days of recovery. Mild damage increased to approximately 30% after stimulation and persisted during early recovery before returning to baseline (Fig. 2F-J, P, and Q). A similar progression was observed following TrpA1 activation: mild damage increased from approximately 5% to 35% immediately after stimulation and returned to baseline by day 7, whereas severe damage increased from approximately 5% to 50% and progressively declined during recovery (Fig. 2A-E, R, and S).

**Figure 2.**
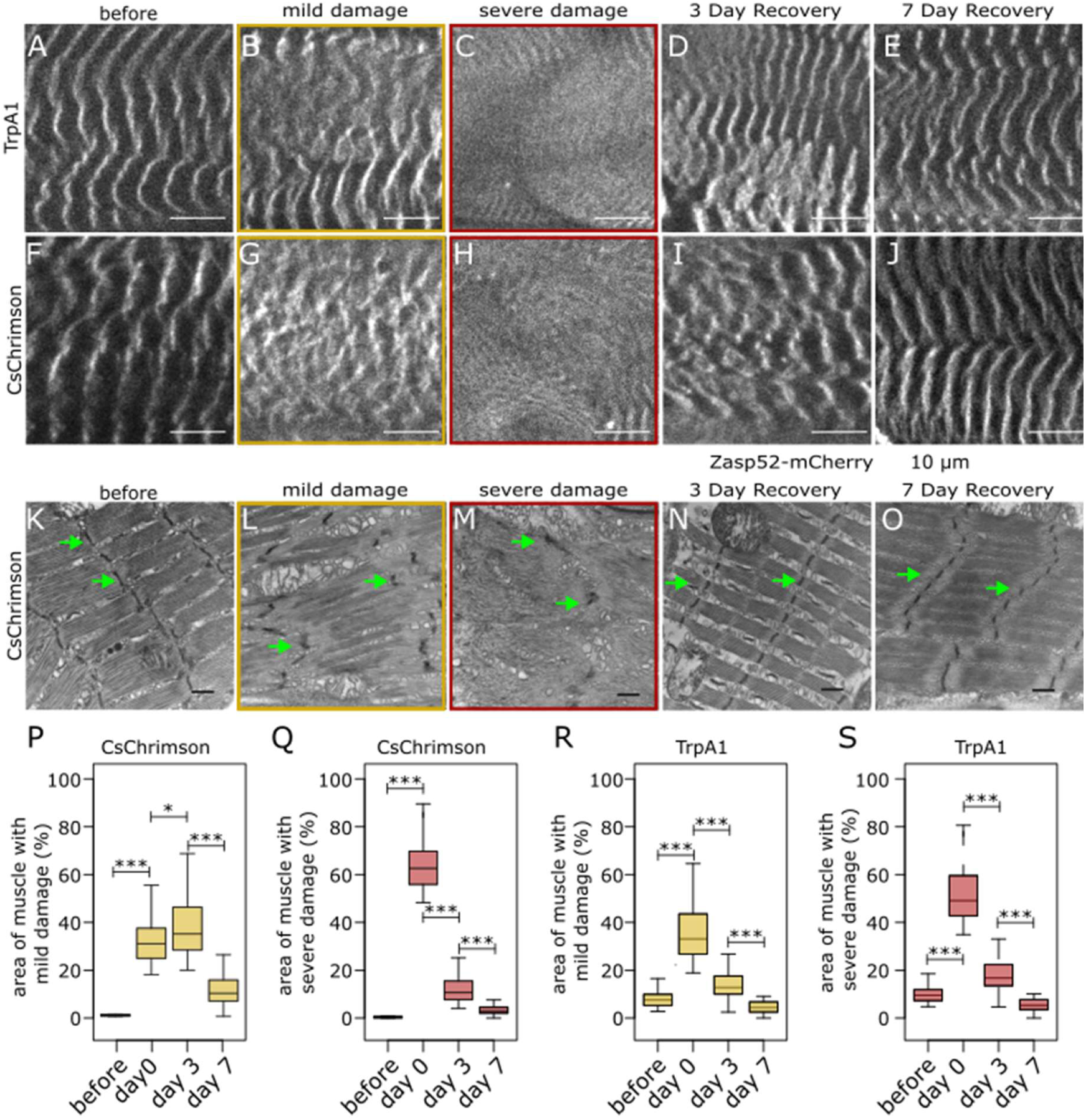
Prolonged depolarization induces massive myofibril disassembly followed by myofibril remodeling. A–E) Confocal images of Zasp52-mCherry-expressing Mef2>TrpA1 levator muscles before depolarization-induced damage (A), immediately after prolonged depolarization (B, C), and 3 and 7 days after depolarization (D, E). B) Representative example of a mildly damaged area. C) Representative example of a severely damaged area. D, E) Representative images of levator muscles 3 and 7 days after depolarization, respectively. E) Sarcomere structure is completely restored by 7 days after depolarization. F–J) Confocal images of Zasp52-mCherry-expressing Mef2>CsChrimson levator muscles before depolarization-induced damage (F), immediately after prolonged depolarization (G, H), and 3 and 7 days after depolarization (I, J). G) Representative example of a mildly damaged area. H) Representative example of a severely damaged area. I, J) Representative images of levator muscles 3 and 7 days after depolarization, respectively. J) Sarcomere structure is completely restored by 7 days after depolarization. K–O) Transmission electron microscopy images of Mef2>CsChrimson levator muscles before depolarization-induced damage (K), immediately after prolonged depolarization (L, M), and 3 and 7 days after depolarization (N, O). L) Representative example of a mildly damaged area. M) Representative example of a severely damaged area. N, O) Representative images of levator muscles 3 and 7 days after depolarization, respectively. O) Sarcomere structure is completely restored by 7 days after depolarization. P–S) Boxplots showing the percentage of muscle area exhibiting damage under different conditions. P) Mild damage following prolonged depolarization using Mef2>CsChrimson reaches approximately 30% of the muscle area immediately after depolarization and decreases to approximately 10% by 7 days. Q) Severe damage following prolonged depolarization using Mef2>CsChrimson reaches approximately 60% immediately after depolarization, decreases rapidly by 3 days, and is completely resolved by 7 days. R) Mild damage induced by Mef2>TrpA1 reaches approximately 30% immediately after prolonged depolarization and is completely resolved by 7 days. S) Severe damage induced by Mef2>TrpA1 reaches approximately 50% immediately after prolonged depolarization and is completely resolved by 7 days. Scale bars: A–J, 10 µm; K–O, 1 µm. Statistical significance was determined using an unpaired two-tailed Welch’s t-test. ns, not significant; P < 0.05; P < 0.01; P < 0.001.

Transmission electron microscopy confirmed extensive structural disruption. Before stimulation, femur muscles displayed the characteristic organization of tubular muscle, with densely packed myofibrils and clearly defined sarcomeres (Fig. 2 K). After 2 hrs of stimulation, myofibrils were largely absent and the remaining Z-discs were markedly reduced. Z-disc spacing increased approximately fivefold, from 191 nm before stimulation to 897 nm after stimulation. By three days, myofibril organization was substantially restored, and by seven days Z-disc spacing had returned to 187–191 nm, indistinguishable from control muscles (Fig. 2K-O and Supplementary Fig. 2D).

### Filamin mechanosignaling regulates myofibril remodeling

Since filamin accumulates at sites of muscle damage and has been proposed to function as a mechanosensor in muscle (Fisher et al., 2024; Leber et al., 2016; Orfanos et al., 2016), we asked whether its mechanosensitive conformational state regulates its localization during myofibril remodeling. We analyzed three GFP-tagged filamin alleles: *wildtype-GFP*, constitutively *closed-GFP*, and constitutively *open-GFP* in combination with the depolarization damage induced by *Mef2>TrpA1*.

Before depolarization damage, wildtype-GFP and closed-GFP localized predominantly to the Z-discs but also showed diffuse cytoplasmic localization, whereas open-GFP is restricted to the Z-discs (Fig. 3A, E, and I). Following TrpA1-induced damage, wildtype-GFP diffused from the Z-discs into prominent cytoplasmic accumulations that were frequently associated with damaged regions (Fig. 3B). Diffusion was reversed during recovery (Fig 3C, D). Closed-GFP showed a similar increase in cytoplasmic accumulation after damage (Fig. 3F-H). Closed-GFP was less often localized to the Z-disc compared to wildtype-GFP before damage (Fig. 3N). Open-GFP remained at the Z-discs throughout damage and recovery (Fig. 3J-L, and N).

**Figure 3.**
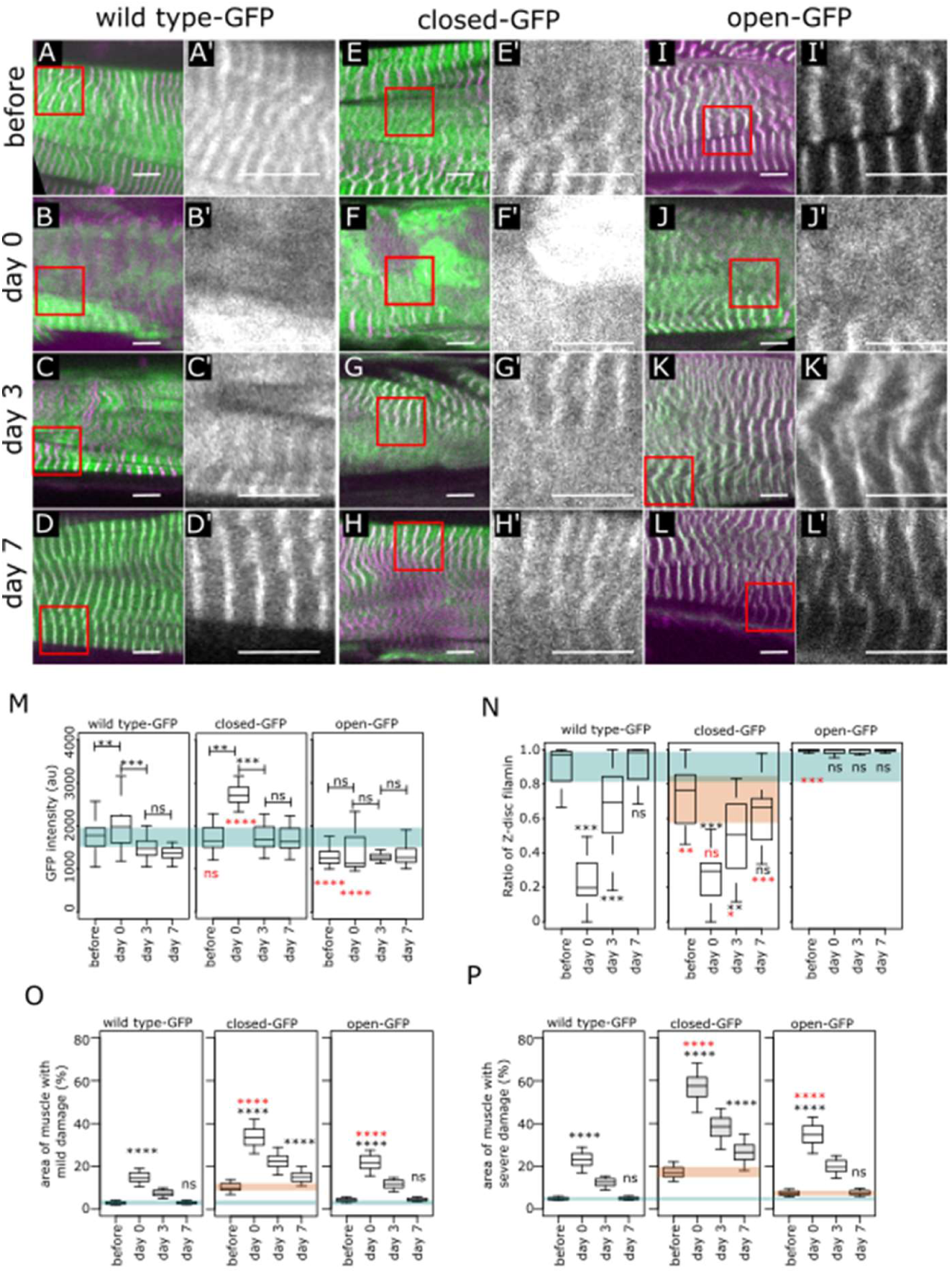
Filamin localization changes following depolarization-induced damage depends on mechanosignaling. A–L) Representative confocal images of levator muscles expressing Zasp52-mCherry, Mef2-Gal4 > UAS-TrpA1, and fluorescently tagged filamin alleles (wild-type-GFP, closed-GFP, or open-GFP). GFP is shown in green and Zasp52-mCherry in magenta. A–D) Representative images from the wild-type-GFP depolarization damage experiment. A) Before depolarization, wild-type-GFP predominantly localizes to the Z-disc, with additional localization at the M-line and diffuse cytoplasmic signal. B) Following depolarization-induced damage, wild-type-GFP forms large cytoplasmic accumulations. C, D) By 3 and 7 days after damage, respectively, wild-type-GFP progressively returns to the Z-disc. E–H) Representative images from the closed-GFP depolarization damage experiment. E) Before damage, closed-GFP predominantly localizes to the Z-disc, with additional M-line and diffuse cytoplasmic localization. F) Following depolarization-induced damage, closed-GFP forms large cytoplasmic accumulations. G, H) At 3 and 7 days after damage, respectively, closed-GFP progressively returns to the Z-disc but does not completely recover its pre-damage localization. I–L) Representative images from the open-GFP depolarization damage experiment. I) Before damage, open-GFP is strictly localized to the Z-disc, with no detectable M-line or diffuse cytoplasmic localization. J) Following depolarization-induced damage, open-GFP remains exclusively localized to the Z-disc, with no formation of large cytoplasmic accumulations. K, L) At 3 and 7 days after damage, respectively, open-GFP remains exclusively localized to the Z-disc. M) Boxplots showing filamin fluorescence intensity in levator muscles before and after depolarization-induced damage. Wild-type-GFP fluorescence intensity increases slightly following damage and returns toward pre-damage levels during recovery. Closed-GFP fluorescence intensity increases substantially following damage and subsequently returns toward pre-damage levels. Open-GFP fluorescence intensity remains largely unchanged following damage. N) Boxplots showing the proportion of muscle area exhibiting Z-disc localization of filamin. Randomly selected positions within the muscle were scored for the presence or absence of wild-type-GFP, closed-GFP, or open-GFP at the Z-disc. Closed-GFP exhibits less Z-disc localization than wild-type-GFP under baseline conditions. Open-GFP is consistently localized to the Z-disc. Following depolarization-induced damage, both wild-type-GFP and closed-GFP show a loss of Z-disc localization. Wild-type-GFP progressively recovers Z-disc localization and returns to pre-damage levels by 7 days, whereas closed-GFP does not fully recover. O) Boxplots showing the percentage of muscle area exhibiting mild damage. Closed-GFP muscles exhibit more mild damage than wild-type-GFP muscles following depolarization. Mild damage in closed-GFP muscles does not fully recover by 7 days, whereas wild-type-GFP muscles return toward baseline. Open-GFP exhibits a damage profile like wild-type-GFP. P) Boxplots showing the percentage of muscle area exhibiting severe damage. Closed-GFP muscles exhibit approximately threefold more severe damage than wild-type-GFP muscles following depolarization and do not fully recover by 7 days. Open-GFP exhibits slightly more severe damage than wild-type-GFP but, like wild-type-GFP, shows complete recovery by 7 days. Scale bar in A–L are 10 µm.

Filamin levels increased following damage in wildtype-GFP and closed-GFP muscles but not in open-GFP muscles (Fig 3N), indicating that this response depends on the ability of the MSR to adopt the closed conformation. Functionally, closed-GFP significantly increased both mild and severe damage and prevented complete recovery (Fig 3O and P), whereas open-GFP produced a more modest increase in damage immediately after stimulation but did not hinder recovery (Fig 3O and P). These results indicate that filamin mechanosignaling and its diffusion from the Z-disc are important for myofibril remodeling.

### Myofibril damage and filamin redistribution increase during aging

We next asked whether filamin mediated muscle remodelling occurs during normal aging. We monitored filamin levels and localization in flies carrying Zasp52-mCherry together with one copy of *wildtype-GFP, closed-GFP, open-GFP,* or *ΔMSR-GFP* (Fig. 4A-F and Supplementary Fig. 3). Wildtype-GFP levels progressively increased during the four weeks of adulthood, with a greater increase in closed-GFP and a smaller increase in open-GFP (Fig. 4G). Consistent with these changes, the proportion of wildtype-GFP localized at the Z-disc decreased with age, accompanied by increased cytoplasmic accumulation (Fig. 4A, B, and H). Closed-GFP showed a similar age-dependent diffusion but had a lower proportion of Z-disc localization at baseline (Fig. 4C, D, and H), whereas open-GFP remained exclusively localized to the Z-disc throughout aging (Fig. 4E, F, and H).

**Figure 4.**
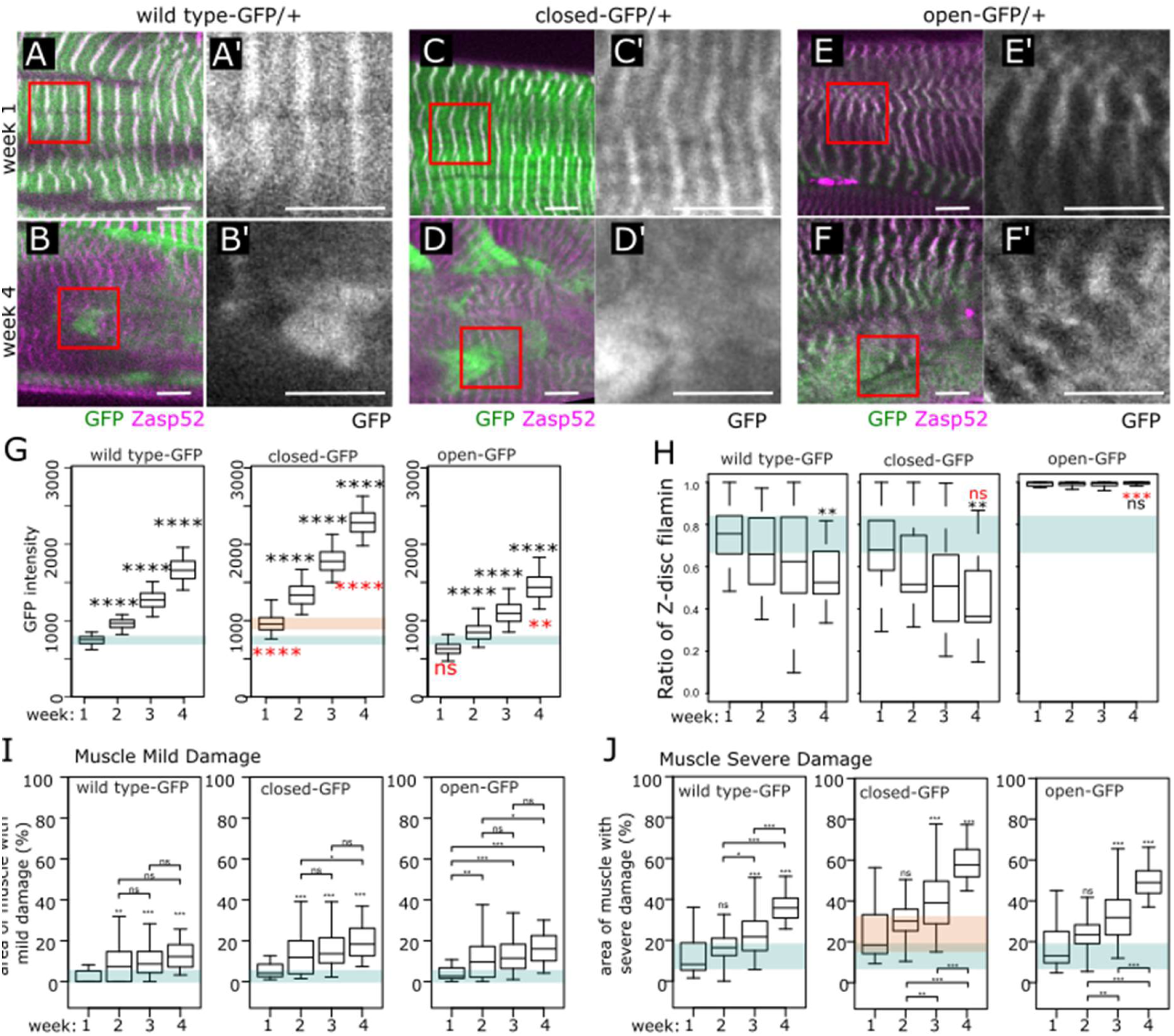
Aging muscles in heterozygous filamin-GFP alleles exhibit progressive filamin accumulation and muscle damage. A–F) Representative confocal images of levator muscles expressing Zasp52-mCherry and fluorescently tagged filamin-GFP alleles at 1 and 4 weeks of age. A) Wild-type-GFP predominantly localizes to the Z-disc in young (1-week-old) flies. B) In 4-week-old flies, wild-type-GFP is predominantly detected in large cytoplasmic accumulations and is associated with areas of muscle damage. C) Closed-GFP predominantly localizes to the Z-disc in young flies. D) In 4-week-old flies, closed-GFP is predominantly detected in large cytoplasmic accumulations and is associated with areas of muscle damage. E) Open-GFP is predominantly localized to the Z-disc in young flies. F) Open-GFP remains predominantly localized to the Z-disc in 4-week-old flies. B, D, F) Representative images of 4-week-old muscles showing mild and severe damage areas. G) Boxplots showing filamin abundance, measured by GFP fluorescence intensity, in different filamin-GFP alleles at 1 and 4 weeks of age. Wild-type-GFP, closed-GFP, and open-GFP fluorescence intensity increases with age, with a greater increase observed in closed-GFP compared with wild-type-GFP. The ΔMSR-GFP allele prevents the age-dependent increase in filamin abundance. H) Boxplots of the proportion of muscle area exhibiting Z-disc localization of filamin. Z-disc localization decreases with age in wild-type-GFP and closed-GFP muscles but is maintained in open-GFP muscles. I) Boxplots showing the percentage of muscle area exhibiting mild damage. All filamin-GFP alleles show a modest age-dependent increase in mild damage. J) Boxplots showing the percentage of muscle area exhibiting severe damage. All genotypes show a substantial increase in severe muscle damage between 1 and 4 weeks of age. Closed-GFP exhibits significantly more severe damage than wild-type-GFP at both 1 and 4 weeks of age, whereas open-GFP exhibits a damage profile like wild-type-GFP. Scale bar in A–F are 10 µm. Statistical significance was determined using an unpaired two-tailed Welch’s t-test. ns, not significant; P < 0.05; P < 0.01; P < 0.001. Black p values are compared to week 1, red is compared to wildtype-GFP.

We next measured age-dependent muscle damage. In *wildtype-GFP* heterozygotes, mild damage increased with age starting at week 2, while severe damage first became apparent at approximately week 3 and increased with age (Fig. 4I and J). *Closed-GFP* heterozygotes exhibited greater severe damage than *wildtype-GFP* but similar amounts of mild damage (Fig. 4I and J). *Open-GFP* did not significantly affect the amount of damage (Fig. 4I and J). These findings suggest that mild myofibrillar damage represents an early stage of age-dependent muscle deterioration that precedes the accumulation of severe damage and that filamin mechanical activation is required for muscle maintenance.

### Homozygous filamin mutants exacerbate age-dependent filamin redistribution

To determine the full consequences of altering the filamin MSR, we examined homozygous knock-in flies. *Wildtype-GFP* homozygotes displayed predominantly Z-disc localization at one week of age but progressively accumulated filamin outside the Z-discs by week 4 (Fig. 5A, B, and H). *Closed-GFP* homozygotes showed cytoplasmic accumulations as early as week 1, which became prominent by week 4 (Fig. 5C, D, and H). In contrast, open-GFP remained localized to the Z-discs at both ages (Fig. 5E, F, and H). Filamin levels increased with age in *wildtype-GFP* homozygotes and were further elevated in *closed-GFP* homozygotes, whereas *open-GFP* levels did not increase with age (Fig. 5G). ΔMSR-GFP levels were too low to accurately measure (Supplementary Fig. 3). A progressive loss of muscle integrity (birefringence) was observed in *wildtype-GFP* (Fig 5I). The filamin mutants at week 4 showed more birefringence loss compared to the *wildtype-GFP* (Fig 5J and Supplementary Fig. 4). These results further support a model in which the filamin MSR controls both the age-dependent accumulation of filamin its localization, and its ability to maintain muscle integrity.

**Figure 5.**
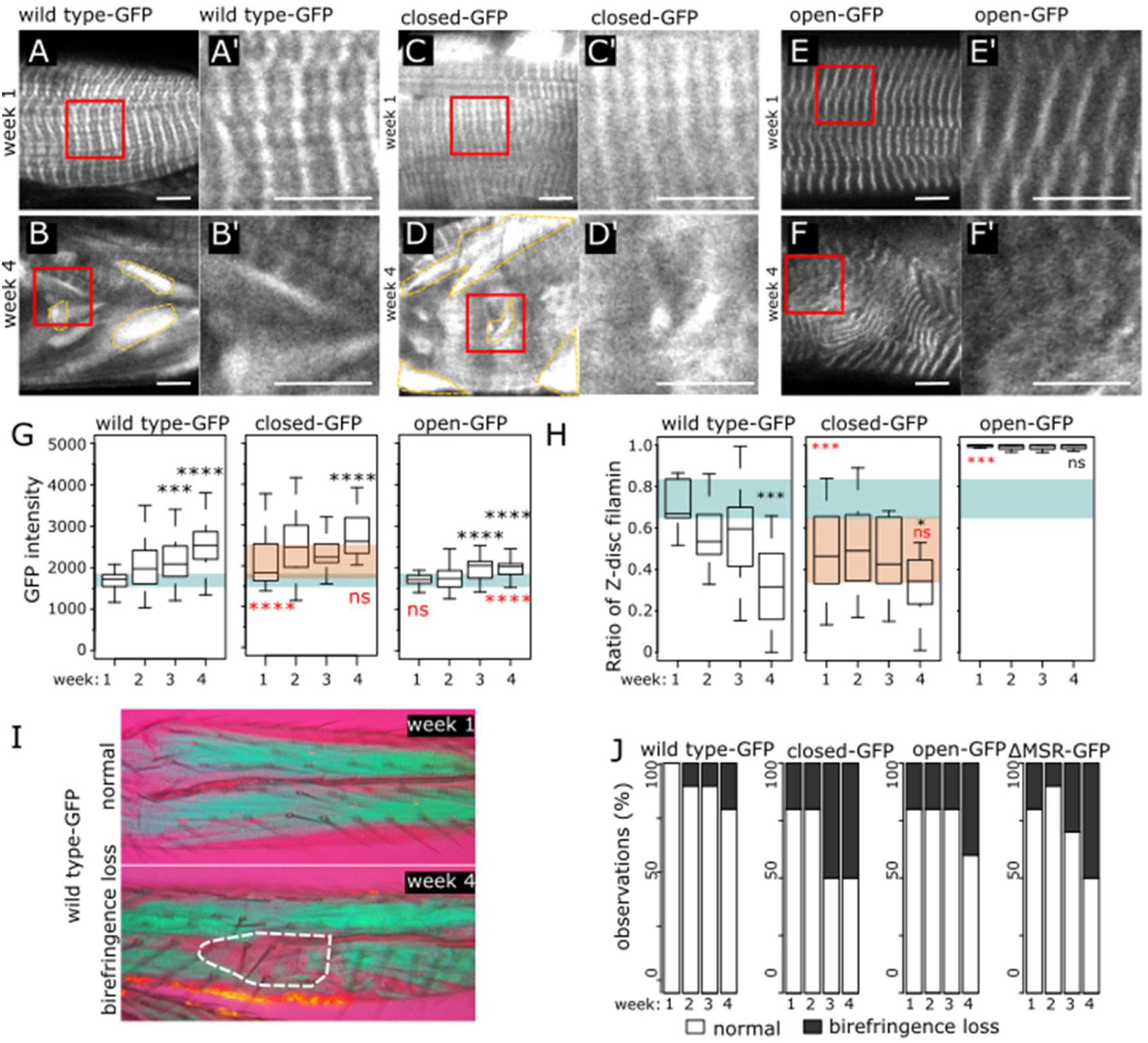
Filamin homozygous mutants reveal mechanisms of mechanosignaling during muscle aging. A–F) Representative confocal images of levator muscles from flies homozygous for different filamin-GFP alleles at 1 week (A, C, E) and 4 weeks (B, D, F) of age. GFP is shown in green. Scale bar in A–F are 10 µm. A) Wild-type-GFP predominantly localizes to the Z-disc at 1 week of age. B) At 4 weeks, wild-type-GFP is predominantly detected in large cytoplasmic accumulations. C) Closed-GFP predominantly localizes to the Z-disc, with some cytoplasmic accumulations, at 1 week of age. D) At 4 weeks, closed-GFP is predominantly detected in large cytoplasmic accumulations. E) Open-GFP is exclusively localized to the Z-disc at 1 week of age. F) Open-GFP remains exclusively localized to the Z-disc at 4 weeks of age; however, wavy myofibrillar patterns indicate extensive muscle damage. G) Boxplots showing filamin abundance, measured by GFP fluorescence intensity, in wild-type-GFP, closed-GFP, and open-GFP homozygous flies at 1 and 4 weeks of age. Wild-type-GFP and closed-GFP fluorescence intensity increases substantially with age, whereas open-GFP fluorescence intensity shows only a minimal increase. H) Boxplots showing the proportion of muscle area exhibiting Z-disc localization of filamin. Wild-type-GFP Z-disc localization decreases with age. Closed-GFP shows lower Z-disc localization than wild-type-GFP, whereas open-GFP remains consistently localized to the Z-disc. I) Polarized light microscopy images of wild-type-GFP legs from 1-and 4-week-old flies. Large areas of reduced birefringence are observed in 4-week-old flies, indicating disruption of muscle structure. J) Quantification of the loss of birefringence in different filamin homozygous genotypes at 1 and 4 weeks of age. Closed-GFP, open-GFP, and ΔMSR-GFP homozygotes exhibit a greater age-dependent loss of birefringence compared with wild-type-GFP homozygotes. Statistical significance was determined using an unpaired two-tailed Welch’s t-test. ns, not significant; P < 0.05; P < 0.01; P < 0.001. Black P values indicate comparisons with 1-week-old flies; red P values indicate comparisons with wild-type-GFP. Scale bar in I are 70 µm.

## Discussion

Our main finding is that femur muscles have a remarkable capacity to repair and remodel their myofibrils following extensive damage. The extent of structural recovery was unexpected given the apparent loss of most myofibrillar structures immediately after stimulation. We show that adult *Drosophila* muscles retain a substantial capacity for myofibril remodelling. We identified two distinguishable states of myofibrillar damage, mild and severe damage. Importantly, our recovery experiments show that even severe myofibrillar damage can be reversed, with myofibrillar organization returning toward a normal state. We further show that muscle damage accumulates with age, with mild damage appearing first and severe damage increasing later in life. This suggests that, like vertebrate muscles, the capacity for effective muscle repair and remodeling declines with age.

Filamin diffusion outside the Z-disc has been observed in vertebrate muscle and in a subset of hereditary myopathies (Chevessier et al., 2015; Orfanos et al., 2016), where it has been used as an indicator of damage. We show that in *Drosophila,* that effective remodeling appears to require transitions between filamin open and closed states. Open filamin remains anchored at the Z-disc and fails to diffuse following damage, whereas closed filamin diffuses but is associated with increased damage and impaired recovery. Thus, the dynamic cycling of filamin between Z-disc-associated and cytoplasmic states are a critical component of the repair response. We propose that mechanical damage induces a conformational cycle in which filamin first adopts an open state associated with damage sensing, followed by a closed state that permits diffusion from the Z-disc.

The accumulation of filamin outside the Z-disc during aging links the damage response to age-dependent muscle deterioration. The progressive diffusion of wildtype filamin with age resembles the response observed following depolarization-induced damage, suggesting that aging muscles experience repeated mechanical damage that progressively exceeds their capacity for repair. The enhanced damage observed in the filamin mutants further suggests filamin mechanosignaling contributes to muscle maintenance.

## Materials and methods

### Drosophila maintenance and stocks

Drosophila stocks were raised and maintained using standard cornmeal media in a 25° C incubator. As control strain we used Oregon Red flies. The *Zasp52^-mCherry^* gene trap was created by replacing the *Zasp52^MI02988^* MIMIC transposon with an artificial exon encoding the *mCherry* sequence (González-Morales et al., 2023). The *cheerio/filamin* replacement alleles *filamin ^WT-GFP^*, *filamin ^open-GFP^*, *filamin ^closed-GFP^*, and *filamin ^ΔMSR-GFP^*were previously described, and all have a C-terminal GFP tag (Huelsmann et al., 2016). The *filamin ^open-GFP^* allele has two amino acid substitution in Ig 16 and Ig18 that prevent the MSR from adopting a closed conformation. The *filamin ^closed-GFP^* allele has a replacement that increases the force required for the MSR to open. The *filamin ^ΔMSR-GFP^* is an in-frame deletion of the Ig domains 14 to19, which compose the MSR region. The *filamin ^WT-GFP^*is the replacement allele with the wild type filamin sequence.

UAS-mCD8GFP is a viable and fertile third chromosome insertion (RRID:BDSC 5130). UAS-IVS-CsChrimson.mVenus is inserted into attP40 (RRID:BDSC 55135). Mef2-Gal4 is a viable and fertile third chromosome insertion (RRID:BDSC 50742). TrpA1 is inserted into attP16 (RRID:BDSC 26263)(Hamada et al., 2008).

### Fluorescence microscopy

Femur muscle dissection and imaging was done as described previously (Guan et al., 2018). Legs were separated from the thorax using fine forceps. Dissection was done in ATP supplemented buffer (Sodium Phosphate Buffer, pH 7.0 20 mM, MgCl₂ 5 mM, EGTA 5 mM), ATP stock solution 7.5 mM, 0.01%, Tween-20 and 50% Glycerol.), then fixed in 4% formaldehyde for 1 hour. Finally, the legs were mounted in Mowiol glycerol mounting media. The femur muscles were imaged using a ThorLabs Laser Scanning Confocal microscope and a 60X Plan Apo 1.42 NA Nikon oil objective. Image analysis was done using Fiji software (Schindelin et al., 2012). GFP fluorescence intensity, muscle damage, and Z-disc localization were quantified using Fiji. GFP intensity was measured as the mean fluorescence intensity within a 200 × 200-pixel square ROI placed within the muscle. Muscle damage was quantified by manually outlining regions of mild and severe damage using the Freehand Selection tool, and the area of each ROI was measured and expressed as a proportion of the total muscle area. Z-disc localization was assessed by dividing the images into semi-randomly selected regions of similar size and scoring them for the presence or absence of a clearly detectable Z-disc pattern.

### Polarized light microscopy

Drosophila legs were dissected and fixed in 4% paraformaldehyde. Fixed legs were mounted between two crossed polarizers and imaged using a 25× objective. A first-order red (1λ) retardation plate was placed between the specimen and the analyzer to enhance visualization of muscle birefringence. The microscope stage was rotated to orient the femur muscles for maximum birefringence, and images were acquired under identical imaging conditions for comparison between genotypes and ages. Areas of reduced or absent birefringence within the femur muscles were scored as regions of loss of muscle integrity.

### Transmission Electron microscopy

Individual femurs were sectioned from dissected legs using needles and fixed for a minimum of 2 hours in 2.5% glutaraldehyde diluted in 0.1 M sodium cacodylate buffer. Following fixation, samples were rinsed three times for at least 10 minutes each in 0.1 M sodium cacodylate buffer. Secondary fixation was performed for 2 hours using 1% osmium tetroxide, after which samples were briefly rinsed with distilled water. Samples were then incubated overnight at 4 °C in 0.25% uranyl acetate. Dehydration was carried out using a graded acetone series: 50% acetone for 10 minutes; 70% acetone for 10 minutes; 95% acetone for 10 minutes; 100% acetone for 10 minutes; and finally dried 100% acetone for 10 minutes. Samples were infiltrated with Epon– Araldite resin using the following sequence: 3:1 ratio of dried 100% acetone to resin for 3 hours, 1:3 ratio of acetone to resin overnight, and 100% resin for 3 hours. Embedding was completed in 100% Epon–Araldite resin and cured in a 60 °C oven for 48 hours. Thin sections (∼100 nm) were cut using a Reichert–Jung Ultracut E ultramicrotome equipped with a diamond knife and collected on 300 mesh copper grids. Grids were stained with 2% aqueous uranyl acetate for 10 minutes, rinsed twice with distilled water (5 minutes each), stained with lead citrate for 4 minutes, and briefly rinsed again with distilled water before air drying. Samples were imaged using a JEOL JEM 1230 transmission electron microscope operating at 80 kV, and digital images were captured with a Hamamatsu ORCA-HR camera.

### Muscle Damage Setup

For optogenetic experiments, Mef2-Gal4, UAS-CsChrimson flies were reared at 20 °C in darkness on media supplemented with 40 ul of 11-cis-retinaldehyde. Raising flies at 25 °C resulted in lethality. One-week-old adult flies were then transferred to a contained area placed directly under an LED light source consisting of 56 LEDs. Flies were exposed to the light for durations ranging from a few minutes to up to 4 hours. After exposure, the lights were turned off, and the flies were allowed to recover at 20 °C in darkness. For thermal activation experiments, Mef2-Gal4, UAS-TrpA1 flies were reared at 20 °C on standard cornmeal media. Then for heat depolarization, the flies were transfer to a preheated 37 °C incubator in empty vials, after 1 hour, the flies were transferred back to 20 °C for recovery.

### Negative geotaxis climbing assay

To measure the climbing speed of individual flies, we used a modified version of the negative geotaxis climbing assay (Rhodenizer et al., 2008). Briefly, single flies were placed inside a clear Drosophila polypropylene vial and capped with a white foam. A gentle tap made the fly drop and instantly climb upwards. We recorded the climbing using a standard camera. Then, we use AnimalTA software to extract the trajectories of individual flies and calculate their moving average velocities (Chiara & Kim, 2023).

